# Revealing the spatiotemporal dynamics of methionine metabolism with a genetically encoded single-fluorophore biosensor

**DOI:** 10.64898/2026.06.05.730515

**Authors:** Katarina A. Cohen, Arnav Jhawar, Gilberto Garcia, Naiara Goni, Megan Chen, Athena Alcala, Ryo Higuchi-Sanabria, Danielle L. Schmitt

## Abstract

Methionine is an essential amino acid, used for protein synthesis, redox homeostasis, and methylation reactions throughout the cell. However, the compartmentalized dynamics of methionine have remained elusive, due to a lack of available tools to measure methionine with high spatial and temporal resolution. To address this limitation, we have developed a single fluorescent protein-based methionine optical reporter (Meteor) which reports subcellular changes in methionine with high dynamic range. Using Meteor, we demonstrate the subcellular uptake of methionine in multiple cell lines into several locations, including the mitochondrial matrix. Furthermore, we use Meteor to illuminate the dynamics of the methionine cycle in the cytoplasm and nucleus, finding cancer cells can rapidly increase methionine from metabolic precursors in both locations. Finally, demonstrated that Meteor can be used to visualize methionine dynamics *in vivo* using *Caenorhabditis elegans*. Thus, we have developed a new tool to measure methionine dynamics across scales with high dynamic range and spatiotemporal resolution.

## Introduction

Methionine is an essential amino acid that is not synthesized *de novo* in humans but is key in metabolic function, as it provides substrates for the folate cycle, methylation reactions, polyamine metabolism, and redox maintenance through the methionine cycle^1,2^. Methionine enters the methionine cycle and is converted to SAM in all mammalian cells and tissues, but the majority of this flux occurs in the liver^3–5^. The alteration of methionine metabolism is a predominant feature in the onset and development of chronic liver diseases^6,7^. The dysregulation of methionine metabolism is also associated with cancer pathogenesis, as many cancers are reported to be dependent upon exogenous methionine for proliferation^8,9^. Cancer cells can synthesize methionine from its metabolic precursor, homocysteine, so dependency of cancer cells on exogenous methionine in the presence of homocysteine highlights altered metabolic flux in methionine-linked pathways^10,11^. Methionine restriction is also reported to increase the lifespan of many model organisms, emphasizing the key role methionine plays in cell and organismal function and survival^12^. Despite the pivotal role methionine plays in a variety of biological contexts, we lack effective tools to study methionine dynamics across scales, from the subcellular to multi-cellular organisms.

Most studies of the methionine cycle focus on cytoplasmic methionine pools, but the nucleus, mitochondrial matrix, and endoplasmic reticulum have been recently identified as potential subcellular locations where methionine is locally consumed^13–16^. However, the compartmentalized uptake and use of methionine remain underexplored because typical methods to measure intracellular methionine, including mass spectrometry, metabolic flux analysis, and *in vivo* imaging using positron emission tomography of ^11^C-methionine, can miss spatiotemporal dynamics at the single cell level^17–20^. When metabolites can be measured in single cells, temporal resolution is often sacrificed^21^. Although these approaches have enabled the investigation of the methionine cycle and tumor imaging in humans, they cannot acquire the real-time visualization of methionine flux with the high spatial and temporal resolution needed to better understand the dynamics of methionine utilization^22^.

Genetically encoded biosensors are well-suited to study compartmentalized metabolism because biosensors can be expressed in living cells to fluorescently report intracellular activity, such as metabolite levels, with spatiotemporal and subcellular resolution^23^. Previous efforts have resulted in biosensors for methionine, but these reporters were limited in their application, not used in mammalian systems, and not used to explore the subcellular dynamics of methionine. Notably, two Förster Resonance Energy Transfer (FRET)-based methionine biosensors have been developed. The first was used to monitor the intracellular methionine level in bacteria and yeast but was limited in dynamic range and multiplexing capacity^24^. The second methionine biosensor incorporated a fluorescent unnatural amino acid (CouA), but its use was never demonstrated in cellular systems^25^.

To overcome these limitations, we developed a single-fluorophore excitation-ratiometric biosensor for methionine termed Meteor (<u>met</u>hionin<u>e</u> <u>o</u>ptical <u>r</u>eporter). We demonstrate the use of Meteor to measure methionine using multiple modalities and across scales. With Meteor, we measured subcellular methionine import in multiple mammalian cell lines, revealing the real-time flux on methionine, and used Meteor to measure the subcellular dynamics of the methionine cycle. Finally, we demonstrate that Meteor reports methionine dynamics *in vivo* in *C. elegans*. Altogether, we present a novel methionine biosensor with large dynamic range to uncover the compartmentalized dynamics of methionine across scales.

## Results

### Development of Meteor

In developing a fluorescent protein-based reporter for methionine, we selected the *N. meningitides* methionine periplasmic binding protein MetQ as the sensing domain (Fig 1a)^26^. Circularly permuted superfolder green fluorescent protein (cpsfEGFP) was inserted into flexible regions of the protein, particularly at sites that experience a conformational change as determined by calculating the change in α dihedral angle (Supplemental Fig 1a). cpsfEGFP was inserted into 30 positions within these regions. N-terminal Ser/Ala/Glu and c-terminal Gly/Thr linkers were affixed between cpsfEGFP and the sensing domain. To assess each candidate biosensor, we measured the response of each candidate to methionine in a clarified bacterial lysate. We measured the change in fluorescence upon methionine addition at 480 nm excitation and 520 nm emission (ΔF/F). Of the candidates, only the biosensor with cpsfEGFP inserted after Val 51 in MetQ had a consistent response with ΔF/F of 0.22 ± 0.016 (Fig 1b). This variant was selected for further optimization.

**Figure 1.**
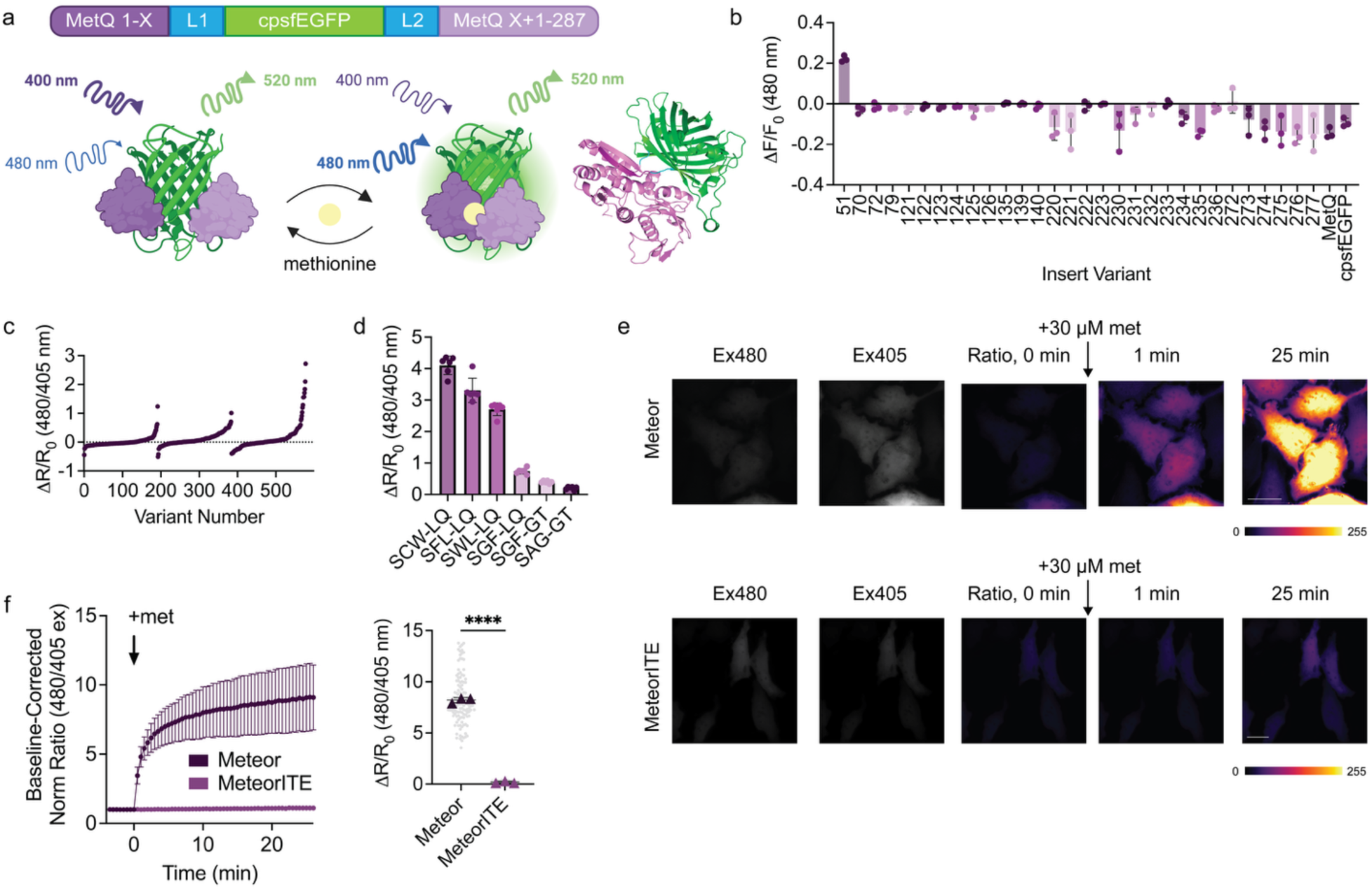
Development of Meteor. a, (top) The design and domain layout of Meteor with cpsfEGFP, bookended by linkers, inserted into MetQ at amino acid residue 51. (bottom left) A cartoon diagram of Meteor that demonstrates its ratiometric increase upon reversibly binding methionine. (bottom right) An Alphafold3 predicted structure of Meteor (green: cpsfEGFP. purple: MetQ, cyan: linkers). Created in BioRender, Cohen, K.A. (2026). b, Fluorescence change (ΔF/F_0_) after addition of 100 µM methionine of initial Meteor variants in clarified bacterial lysate (n = 3 technical replicates). c, Ratio change (ΔR/R_0_) of linker variants screen in clarified bacterial lysate with 100 μM L-methionine across three linker screenings. d, Ratio change of top performers obtained from linker screening in clarified bacterial lysate and treated with 100 μM L-methionine (n = 6 technical replicates). e, (top) Representative pseudocolor images of the ratio response of HeLa cells expressing Meteor treated with 30 µM methionine. (bottom) Fluorescent microscopy images of the ratio response of HeLa cells expressing MeteorITE treated with 30 µM methionine. f, (left) Average response of Meteor and MeteorITE treated with 30 μM methionine addition in HeLa. (right) Maximum ratio change (ΔR/R_0_) of Meteor and MeteorITE in response to 30 μM methionine. Cells are shown in small, gray circles, and the average from each independent experiment is shown in a colored triangle. **** P < 0.0001 by unpaired two-tailed t-test. Meteor n = 121 cells from 3 independent experiments. MeteorITE n = 105 from 3 independent experiments. Scale bar represents 20 μm. For all graphs, mean ± standard deviation is shown.

We conducted three rounds of linker screening to enhance the modest response of the preliminary reporter (Fig 1c). The linkers that connect cpsfEGFP to the sensing domain are crucial for biosensor functionality, with mutations in either linker often resulting in significant changes to the biosensor’s response^27–28^. During our screening, we evolved towards an excitation-ratiometric biosensor, selecting for variants which demonstrated a change in 405 nm excitation-emission and 480 nm excitation-emission (ΔR = 480 nm ex-em/405 nm ex-em). We and others have generated excitation-ratiometric biosensors, which benefit from being generally pH insensitive and measurements are independent of biosensor concentration^29–31^. We generated libraries of linker variants, and 192 variants were randomly selected from each library and tested in clarified bacterial lysate. Three iterations of linker screening produced a variant with Ser/Cys/Trp-Leu/Glu linkers and ΔR/R of 4.11 ± 0.29 in response to 100 μM methionine (Fig 1d). Because this variant successfully reports methionine dynamics with good dynamic range, we selected this top performer for further characterization and termed it Meteor (<u>met</u>hionin<u>e o</u>ptical <u>r</u>eporter).

To further confirm that Meteor’s response is a result of the MetQ sensing domain binding to methionine, we developed MeteorITE (Meteor, <u>it</u>’s binding is <u>e</u>xtinct), a Meteor variant that is unable to bind to methionine. We created singly, doubly, or triply mutated variants, targeting amino acid residues critical to methionine binding in the MetQ binding pocket (Supplement Fig 1b). While these mutations produced suppressed responses compared to Meteor, they still exhibited some level of affinity for methionine or did not express well in mammalian cells. Thus, we randomly mutagenized N238, a residue that forms a crucial hydrogen bond with α-amino and α-carboxyl groups in methionine, and Y81 on the opposite lobe of MetQ that packs around methionine’s thioether group^26^. We screened 192 variants from this library in bacterial lysate and determined which mutants had a similar fluorescent intensity to Meteor while experiencing minimal change upon the addition of methionine (Supplement Fig 1c). The resulting MeteorITE harboring Y81P and N238T expressed well in HeLa cells and exhibited a significantly repressed response to methionine compared to previous nonbinding contenders (Supplement Fig 1d). Thus, we developed a suitable control biosensor.

To determine Meteor’s performance in mammalian systems, Meteor and MeteorITE were expressed in HeLa cells and treated with 30 µM methionine. Upon treatment, we observed an increase in Meteor fluorescence intensity at 488 nm and a decrease at 405 nm indicative of a ratiometric sensor (Supplement Fig 1e). Meteor displayed a ΔR/R_0_ of 8.23 ± 2.37 in response to 30 µM methionine while MeteorITE exhibited minimal response with a ΔR/R_0_ of 0.16 ± 0.13 (Fig 1e-f). Thus, we have developed a new methionine biosensor, Meteor, suitable for measuring methionine dynamics in mammalian cells.

### In Vitro Characterization of Meteor

We then sought to characterize Meteor’s *in vitro* performance. Meteor exhibited a concentration-dependent change in the ratiometric response, reaching a maximum response around 32 mM methionine (Fig 2a, b). Meteor has an affinity for methionine of 1514 µM (95% confidence interval 1459 µM to 1572 µM). The *K*_D_ of MetQ is reported to be approximately 0.2 nM, indicating that the insertion of cpsfEGFP decreased its native affinity for methionine^26^. The concentration of intracellular methionine varies across cell lines and is often estimated to fall within the range of 20 µM to 80 µM with a fast and uniform distribution between intracellular and extracellular methionine pools^32–33^. To determine if Meteor can detect physiological methionine concentrations in unstarved HeLa cells, cell membranes were permeabilized with digitonin, and 1 µM to 500 µM methionine was added to the cells. The average ratio response of Meteor-expressing cells was plotted versus the concentration of methionine to create an in-situ calibration curve. In HeLa cells, Meteor responded robustly to a physiological range of methionine and was determined to be suitable for our experiments (Supplement Fig 1f). Using stopped-flow at a saturating concentration of methionine, we determined the *k_ob_*_s_ of Meteor to be 0.0521 s^-1^ (Supplement Fig 1g).

**Figure 2.**
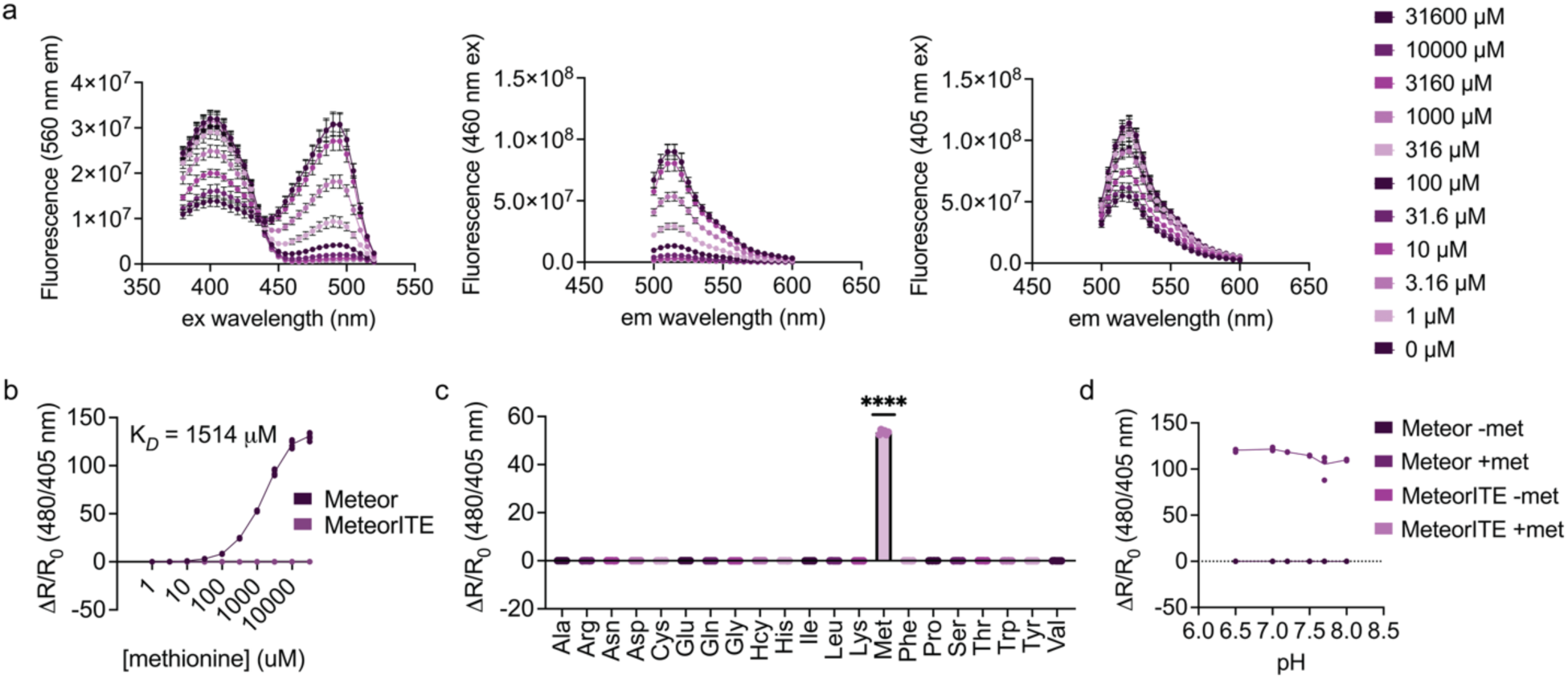
*In vitro* characterization of Meteor. a, (left) Meteor emission sweep from 500-600 nm with excitation at 460 nm in the presence of various concentrations of l-methionine. (middle) Meteor emission sweep from 500-600 nm with excitation at 405 nm in the presence of various concentrations of methionine. (right) Meteor excitation sweep from 360-520 nm excitation with fluorescence measured at 560 nm emission in the presence of various concentrations of methionine. Data represents 1 protein preparation with 3 technical replicates. b, Dose-response curve for Meteor and MeteorITE ratio changes (ΔR/R_0_) in response to multiple concentrations of methionine. Shown are the mean ± standard deviation of 6 technical replications for 2 independent protein preparations of Meteor or MeteorITE. K*_D_* was determined using nonlinear fit to y = B_max_*x(K*_D_* + x). c, Maximum ratio change of Meteor in response to 1 mM of each respective amino acid (n = 6 replicates from 2 independent protein preparations). **** P < 0.0001 by ordinary one-way ANOVA. d, pH dependency of Meteor and MeteorITE in the presence of 0 mM (Meteor dark purple, MeteorITE dark pink; n = 6 from 2 individual protein preparations) or 31.6 mM methionine (Meteor light purple, MeteorITE light pink; n = 6 from 2 individual protein preparations). For all graphs, mean ± standard deviation is shown.

Meteor was also highly specific for methionine, exhibiting a ΔR/R_0_ of 53.39 in response to 1 mM methionine. (Fig 2c). Meteor minimally responded to all other amino acids and the related molecule homocysteine (hcy) at the same concentration. We next assessed the pH-dependency of Meteor. Fluorescent protein-based biosensors are often pH dependent, but ratiometric measurements can produce a biosensor with decreased pH dependency^34–35^. Meteor exhibited a consistent ΔR/R_0_ from pH 6.5 to 8 (Fig 2d). We also found that MeteorITE similarly does not have a change in ΔR/R_0_ based on pH. Thus, Meteor enables the sensitive and specific detection of methionine, and the ratio change (ΔR/R_0_) is consistent across a range of physiological pH.

### Meteor Enables Multi-Modal Detection of Methionine Uptake

With a high dynamic range, we anticipated Meteor would enable multi-modal measurement of methionine dynamics in living cells, which would be suitable across multiple applications. To demonstrate the utility of Meteor for multiple modalities, we first imaged methionine dynamics in HeLa cell populations using a plate reader. Consistent with single-cell data, Meteor had a robust, concentration-dependent increase in response to methionine addition as compared to MeteorITE (Meteor cpsfEGFP/ExRai ratio for 10 µM met = 1.951 ± 0.435; for 100 µM met = 3.163 ± 0.536; for 1 mM met = 3.848 ± 0.624; Fig 3a). Next, we expected that the large increase in 480 nm excitation-emission alone by Meteor would enable robust detection of methionine uptake in cells as measured with flow cytometry. To normalize for biosensor expression levels, we generated a variant of Meteor and MeteorITE fused to the red fluorescent protein mScarlet (Meteor-mScarlet and MeteorITE-mScarlet, respectively; Fig 3b). HeLa cells expressing Meteor-mScarlet or MeteorITE-mScarlet were treated with increasing concentrations of methionine, and flow sorted. Consistent with measurements taken using both fluorescence microscopy and a plate reader, there was a concentration-dependent increase in Meteor:mScarlet ratio that was not seen with MeteorITE:mScarlet ratio (Meteor:mScarlet for 30 µM met = 1.074 ± 0.067; for 300 µM met = 1.421 ± 0.077; Fig 3c-e). Thus, Meteor can be used to measure intracellular methionine dynamics across multiple modalities, enabling broad use of Meteor for methionine sensing in many applications.

**Figure 3.**
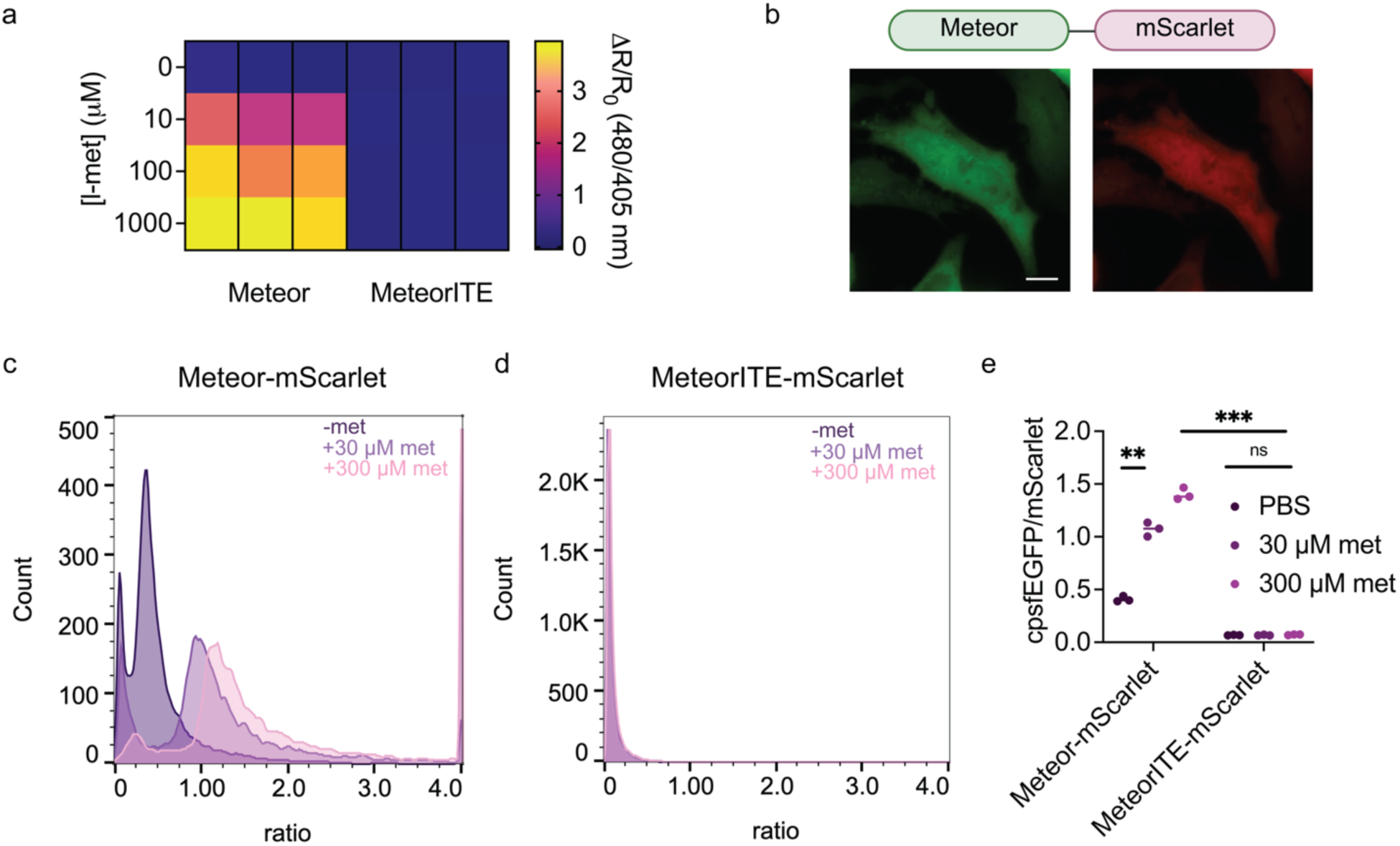
Meteor enables multi-modal detection of methionine uptake. a, Heatmap of excitation-ratio of HeLa cells expressing Meteor and MeteorITE measured using a plate reader. Cells were treated with either a vehicle control or 10, 100, or 1000 μM methionine. The response after 30 minutes was normalized to the response prior to addition. Warmer colors represent higher ratios and more methionine uptake. Data are averages from three independent experiments. b, Representative images of Meteor fused to mScarlet (Meteor-mScarlet) expressed in HeLa cells. Scale bar represents 10 μm. c-d, Representative histograms of the ratio (cpsfEGFP/mScarlet) of HeLa cells in suspension expressing Meteor-mScarlet (c) and MeteorITE-mScarlet (d) with no methionine (dark purple), 30 μM methionine (light purple), or 300 μM methionine (pink). Cells were gated using mScarlet as an expression marker. e, Mean cpsfEGFP/mScarlet ratios of Meteor-mScarlet and MeteorITE-mScarlet HeLa cells measured by flow cytometry with no methionine (dark purple), 30 μM methionine (light purple), or 300 μM methionine (pink) across three biological replicates. Meteor 30 μM vs. PBS ** P = 0.0019; Meteor 300 μM vs. MeteorITE 300 μM *** P = 0.0006; MeteorITE 300 μM vs. PBS n.s. P = 0.2246 by two-tailed paired t-test.

### Measurement of Subcellular Methionine Uptake with Meteor

To investigate the dynamics of compartmentalized methionine uptake, we used targeting motifs to localize Meteor to the cytoplasm, outer endoplasmic reticulum, mitochondrial matrix, and nucleus, and expressed in HeLa cells (Fig 4a-d). While the methionine cycle is considered largely cytosolic, methionine adenosyltransferase1/2 (MAT1/2) isoenzymes and MAT2 subunits have been detected in the nucleus, and the MAT1α has been found in the mitochondrial matrix^36–39^. Thus, these locations were selected because methionine is known to be used by MATs at these locations, or these were previously identified locations for local methionine use. When HeLa cells were treated with methionine, we observed robust methionine uptake at all measured subcellular locations (Meteor ΔR/R: cyto = 15.00 ± 4.26; ER = 8.41 ± 2.93; mitochondrial matrix = 1.80 ± 0.90; nucleus = 6.87 ± 2.65; Fig 4e-f). As a metric for the rate of methionine import, we also calculated the time-to-half maximum (t_1/2_) of Meteor response and found methionine import to all locations had similar kinetics (Meteor t_1/2_: cyto = 1.67 ± 0.60 min; ER = 1.73 ± 1.07 min; mitochondrial matrix = 1.14 ± 0.32 min; nucleus = 1.72 ± 0.54 min; Fig 4g).

**Figure 4.**
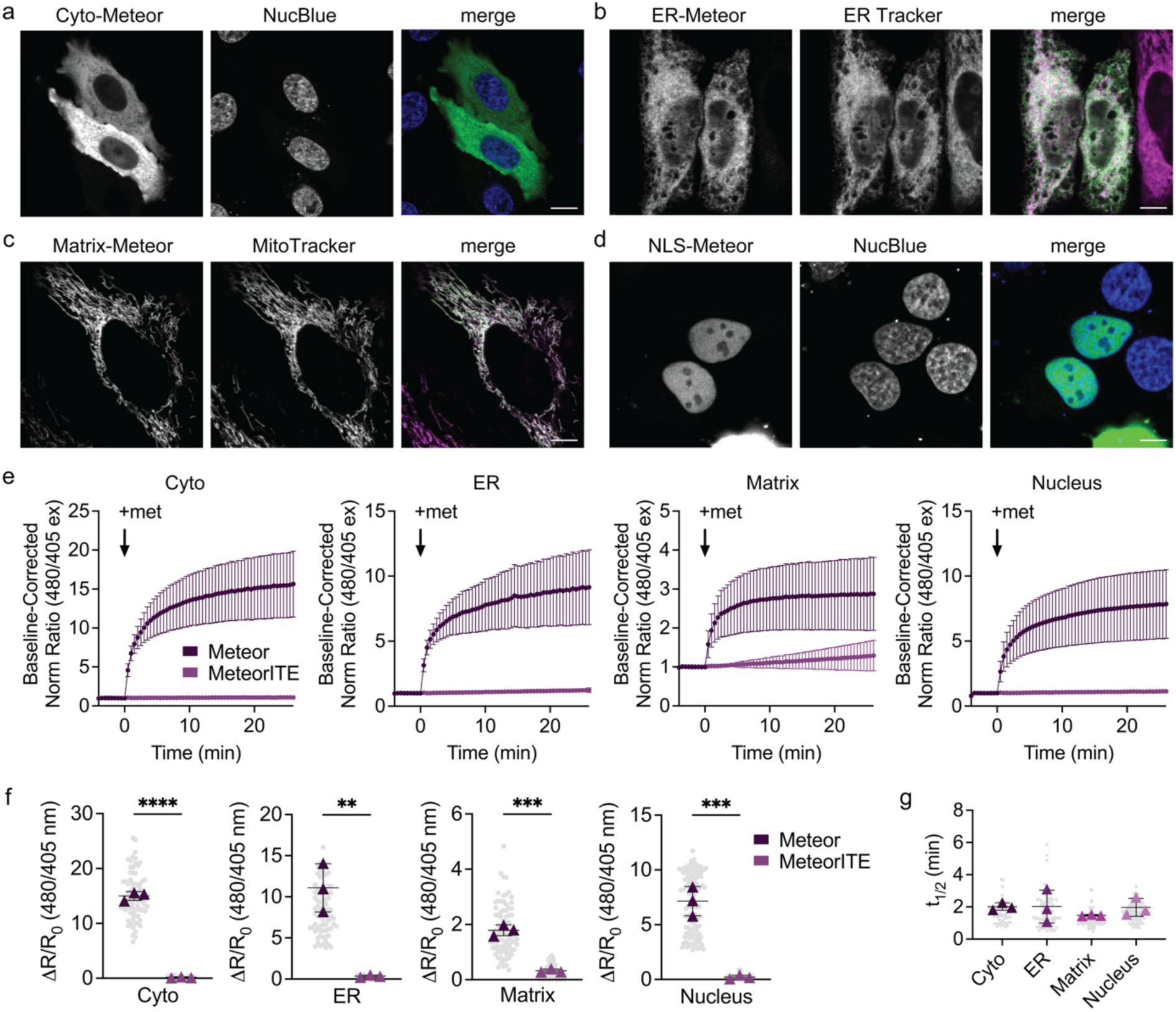
Measurement of subcellular methionine uptake with Meteor. a-d, Representative images of HeLa cells expressing subcellular targeted Meteor to the cytoplasm, outer endoplasmic reticulum, mitochondrial matrix, and nucleus stained with indicated organelle marker. Scale bar represents 10 μm. e, Normalized response of localized Meteor (dark purple) and MeteorITE (light purple) to 30 μM methionine addition in HeLa cells. From left to right, Meteor and MeteorITE are localized to the cytoplasm, outer endoplasmic reticulum, mitochondrial matrix, and nucleus. f, From left to right, the average maximum ratio change of Meteor (dark purple) and MeteorITE (light purple) in response to 30 μM methionine at the cytoplasm, outer endoplasmic reticulum, mitochondrial matrix, and nucleus. Cyto **** P < 0.0001; ER ** P = 0.0031; matrix *** P = 0.0003; nucleus *** P = 0.0009 by unpaired two-tailed t-test. Cyto-Meteor n = 73 cells from 3 independent experiments. Cyto-MeteorITE n = 74 from 3 independent experiments. ER-Meteor n = 75 cells from 3 independent experiments. ER-MeteorITE n = 72 from 3 independent experiments. Matrix-Meteor n = 81 cells from 3 independent experiments. Matrix-MeteorITE n = 86 from 3 independent experiments. NLS-Meteor n = 110 cells from 3 independent experiments. NLS-MeteorITE n = 90 from 3 independent experiments. g, Time-to-half-maximum (t1/2) of subcellular targeted Meteor response to 30 μM methionine from experiments shown in Fig 4e-f. n.s. P = 0.6297 by ordinary one-way ANOVA; pairwise comparisons are all n.s. P > 0.05 by Tukey’s multiple comparisons test. For all graphs, individual cells are shown in small, gray circles, and the average of each independent experiment is shown in a colored triangle.

We next wanted to evaluate both the performance of Meteor and characterize methionine dynamics in multiple cell types. Hepatic models are especially relevant as hepatocellular methionine cycle enzymes are highly active and tightly regulate SAM production from methionine^40^. Localized Meteor variants and corresponding MeteorITE variants were expressed in HepG2 hepatocellular carcinoma cells, IHH immortalized human hepatocytes, and U2OS osteosarcoma cells. Robust methionine uptake was observed at all locations across all cell types (Supplement Fig 2a-c). Like HeLa cells, these cell types displayed similar rates of methionine import across monitored cellular compartments (Supplement Fig 2d-f).

### Meteor Reveals Compartmentalized, Real-Time Dynamics of the Methionine Cycle

The methionine cycle is primarily thought to occur in the cytoplasm, and little is known about the compartmentalized regulation of methionine metabolism, and if cytoplasmic flux of the methionine cycle influences other compartments. We hypothesized that Meteor would allow for the subcellular dissection of methionine dynamics. We focused on methionine production in the cytoplasm or nucleus of HeLa, HepG2, and IHH cells, cell lines which display some degree of methionine dependence. In addition to the cytoplasm, we measured methionine dynamics in the nucleus because there is emerging evidence of methionine metabolism localized to the nucleus^36–38^. First, we examined the activity of methionine synthase, a methionine cycle enzyme that remethylates homocysteine to methionine (Fig 5a). To measure the flux of methionine synthesis, we treated methionine-starved cells expressing Meteor targeted to either the cytoplasm or nucleus with hcy. All cell types readily and rapidly produced methionine from homocysteine, as we observed an increase in Meteor response in the cytoplasm and nucleus compared to MeteorITE control (Meteor ΔR/R: cyto = 0.6912 ± 0.1491; nucleus = 0.6796 ± 0.1885; Fig 5b; Supplemental Fig 3a-c, g-i).

**Figure 5.**
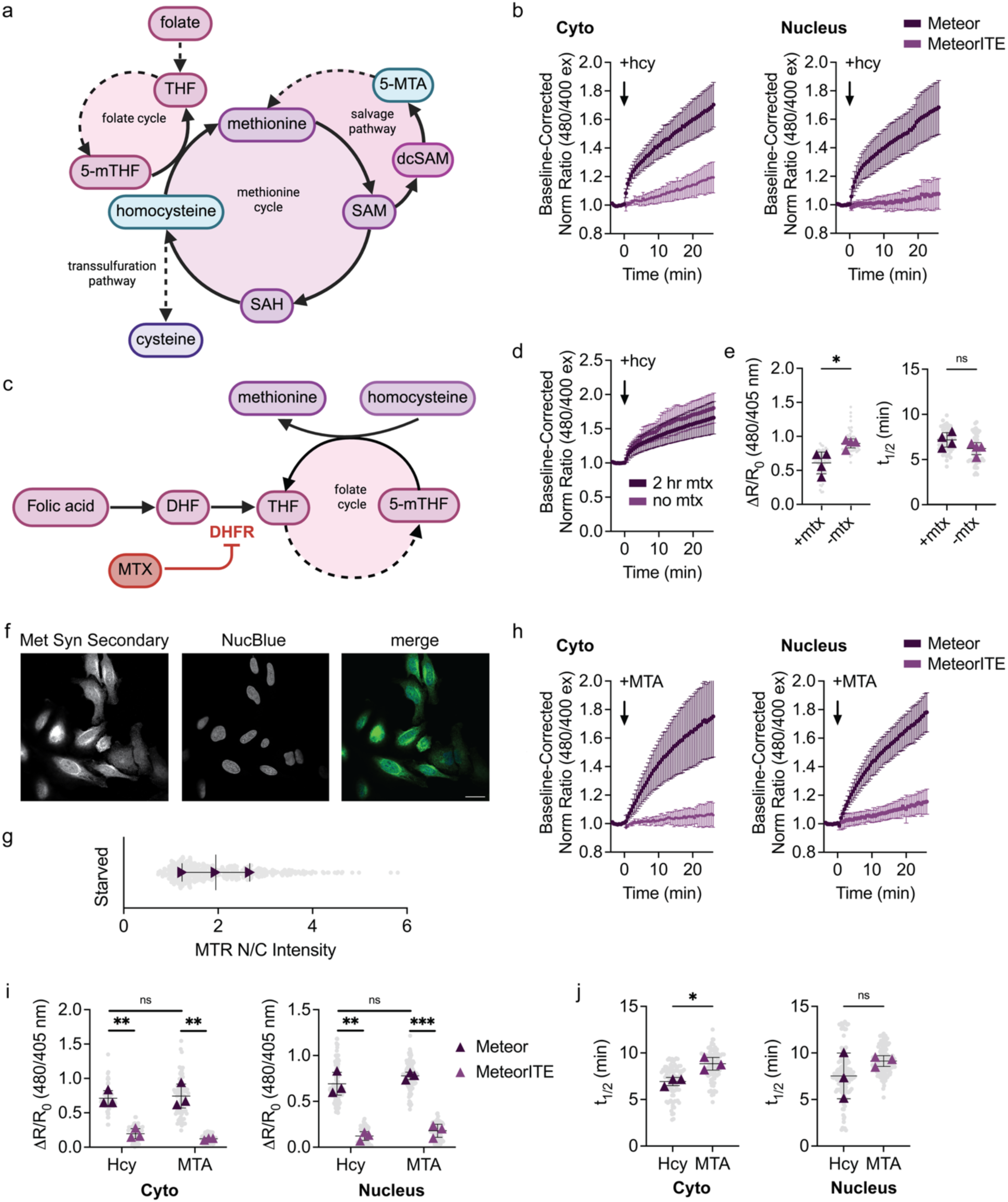
Meteor reveals compartmentalized, real-time dynamics of the methionine cycle. a, Diagram depicting the methionine cycle and interconnection with the methionine salvage pathway and folate cycle. Homocysteine (hcy) and 5’-methylthioadenosine (MTA) are highlighted in blue and were the metabolites used to study methionine synthesis. Created in BioRender, Cohen, K.A. (2026). b, Normalized response of localized Meteor (dark purple) and MeteorITE (light purple) to 150 μM hcy addition in HeLa cells at the (left) cytoplasm and (right) nucleus. c, Diagram depicting the impact of methotrexate (mtx) on the synthesis of methionine from homocysteine. Mtx inhibits the enzyme dihydrofolate reductase (DHFR), depleting 5-methyltetrahydrofolate (5-mTHF) and disrupting the conversion of homocysteine to methionine. Created in BioRender, Cohen, K.A. (2026). d, Normalized response of cyto-Meteor when incubated in normal conditions (light purple) or with 100 μM mtx (dark purple) for 2 hours prior to imaging with 150 μM hcy addition in HeLa cells. e, Maximum ratio change of cyto-Meteor in response to 150 μM hcy in HeLa cells incubated with (dark purple) or without (light purple) 100 μM mtx. * P = 0.0163 by unpaired two-tailed t-test. With mtx n = 63 cells from 4 independent experiments. Without mtx n = 50 cells from 4 independent experiments. f, Representative images of immunostaining for methionine synthase. Scale bar represents 20 μm. g, Nuclear to cytoplasmic ratio of methionine synthase intensity in starved HeLa cells. h, Normalized response of localized Meteor (dark purple) and MeteorITE (light purple) to 30 μM MTA addition in HeLa cells at the (left) cytoplasm and (right) nucleus. i, Maximum ratio change of Meteor and MeteorITE in response to either 150 μM hcy or 30 μM MTA at the (left) cytoplasm and (right) nucleus. Cyto hcy ** P = 0.0024; Cyto MTA ** P = 0.0035; Cyto hcy vs. MTA n.s. P = 0.8030; NLS hcy ** P = 0.0018; NLS MTA *** P = 0.0002; NLS hcy vs. MTA n.s. P = 0.3179 by unpaired two-tailed t-test. Cyto-Meteor hcy n = 97 from 3 independent experiments. Cyto-MeteorITE hcy n = 146 cells from 3 independent experiments. NLS-Meteor hcy n = 110 cells from 3 independent experiments. NLS-MeteorITE hcy n = 124 cells from 3 independent experiments. Cyto-Meteor MTA n = 86 from 3 independent experiments. Cyto-MeteorITE MTA n = 62 cells from 3 independent experiments. NLS-Meteor MTA n = 159 cells from 3 independent experiments. NLS-MeteorITE MTA n = 123 cells from 3 independent experiments. j, Time-to-half-maximum (t1/2) of subcellular targeted Meteor response to 30 μM methionine at the (left) cytoplasm and (right) nucleus in live HeLa cells from experiments shown in Fig 5b-d. Cyto * P = 0.0152; Nucleus n.s P = 0.3347 by unpaired two-tailed t-test.

To convert hcy to methionine, hcy must be methylated by homocysteine methyltransferase, using N^5^-methyl tetrahydrofolate (N^5^-methyl THF). N^5^-methyl THF is produced by the folate cycle, which is inhibited by anticancer drugs including methotrexate (mtx; Fig 5c). To determine the spatiotemporal interplay between the folate cycle and the methionine cycle, we incubated HeLa cells expressing Meteor in the cytoplasm or nucleus with mtx prior to treating with hcy. In mtx-treated cells, we observed suppressed cytoplasmic methionine production, with similar methionine production kinetics to untreated cells (Fig 5d-e, Meteor ΔR/R: without mtx = 0.8992 ± 0.2096; with mtx = 0.5991 ± 0.1898).

Meteor reported similar dynamics and kinetics at both the cytoplasm and nucleus for methionine synthesis from hcy. To determine if this was the result of localized nuclear production, we immunostained for methionine synthase, the enzyme that converts hcy to methionine (Fig 5f). We found consistent expression of methionine synthase throughout the cell with enrichment in the cytoskeleton, consistent with other reports of methionine synthase localization^41^. We found that a small percentage of methionine synthase was nuclear, perhaps indicating localized production (Fig 5g). However, methionine is a small metabolite that could diffuse into the nucleus at a rate faster than can be measured using Meteor.

Similarly, we looked at the methionine salvage pathway, wherein 5’-methylthioadenosine (MTA) is another metabolic precursor to methionine (Fig 5a). Starved cells expressing Meteor targeted to either the cytoplasm or nucleus were treated with MTA. All cell types readily and rapidly produced methionine from MTA, as we observed an increase in Meteor’s response in the cytoplasm and nucleus compared to MeteorITE control (Meteor ΔR/R: cyto = 0.7423 ± 0.2757; nucleus = 0.7799 ± 0.1334; Fig 5h, Supplemental Fig 3d-f, j-l).

Meteor offers the capacity to quantitatively explore the kinetics of methionine synthesis. While the maximum response of Meteor did not significantly change based on the added precursor, in the cytoplasm of HeLa cells, the rate of methionine synthesis from homocysteine was significantly increased compared to MTA (Meteor t_1/2_: cyto hcy = 7.019 ± 1.664 min; cyto mta = 8.737 ± 1.580 min; nucleus hcy = 7.623 ± 2.539 min; nucleus mta = 9.155 ± 1.343 min; Fig 5i-j). Altogether, this data suggests differential rates of methionine production based on the pathway used.

A key enzyme involved in the production of methionine from MTA is methylthioadenosine phosphorylase (MTAP). The homozygous deletion of MTAP is characteristic of many cancer cell lines, including the lung adenocarcinoma cell line A549^42^. To assess how loss of MTAP influences the dynamics of the methionine cycle, we expressed Meteor in the cytoplasm and nucleus of A549 cells. When treated with MTA, A549 minimally produced methionine, as measured by Meteor in the cytoplasm and nucleus (Supplemental Fig 3m). Meanwhile, A549 could produce methionine from homocysteine (Supplemental Fig 3n). Thus, using Meteor, we were able to measure subcellular methionine cycle dynamics in single cells, and determine how oncogenic-associated mutations influence spatiotemporal methionine cycle flux.

### Meteor Measures In Vivo Methionine Dynamics using a C. elegans Model

*Caenorhabditis elegans* is a model system amenable to methionine cycle studies with Meteor because of its highly conserved one-carbon metabolism, rapid life cycle, and transparent body^43–44^. Pharmacological, dietary, or genetic manipulation of the methionine cycle is reported to increase lifespan of *C. elegans*, making this an ideal model organism to study methionine dynamics^45^. We sought to express Meteor in *C. elegans* and monitor methionine dynamics in the intestine upon methionine starvation and methionine refeeding (Fig 6a). *C. elegans* expressing Meteor-mRuby were grown to day 1 and imaged in the absence of food for 270 min (Fig 6b). As *C. elegans* exhibit autofluorescence in blue light (400 nm ex), and we previously demonstrated the change in 480 nm ex-em is sufficient to measure methionine dynamics (Fig 3), we imaged the change in Meteor 480 nm ex-em, normalized to mRuby. In response to starvation, the Meteor response decreased over time, indicating decreased methionine in the gut (Fig 6b). Importantly, the expression of RFP did not change over time, indicating changes observed are due to depletion in methionine and not a reduction in protein levels during starvation (Supplemental Fig 4a). The same cohort of Meteor-RFP-expressing C. elegans were starved overnight and then refed. After refeeding, the Meteor response increased within 90 min, indicating increased intestinal methionine levels (Fig 6c). Again, mRuby levels did not change over time, suggesting protein levels were not changing during refeeding (Supplemental Fig 4b). Thus, we have leveraged Meteor to image the real-time dynamics of methionine in the intestine of *C. elegans*, enabling measurement of methionine dynamics in live, multi-cellular organisms.

**Figure 6.**
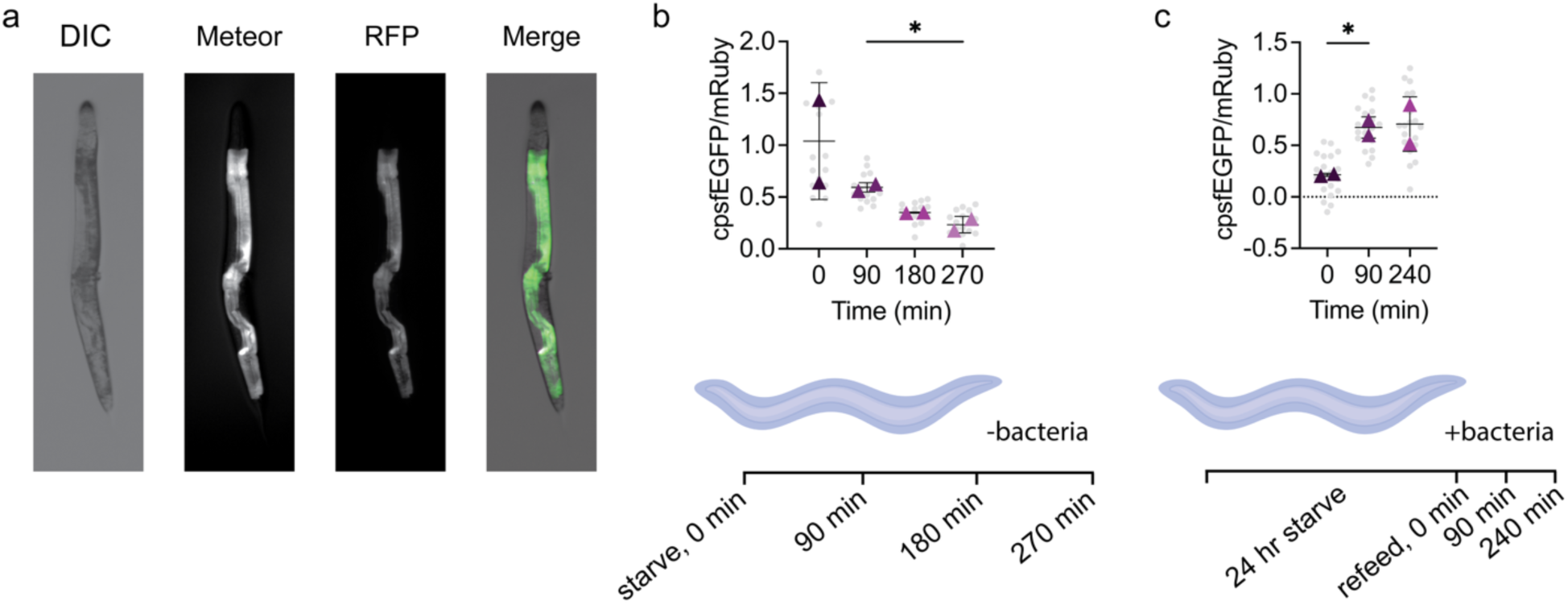
Meteor measures *in vivo* methionine dynamics in a *C. elegans* model. a, Representative images of *C. elegans* expressing Meteor. The pictured *C. elegans* were starved for 24 hours and then refed to monitor Meteor’s response. b, (top) Average ratio response of Meteor across the starvation period. (bottom) Starvation time course that was monitored. 90 min vs. 270 min * P = 0.0244 by unpaired two-tailed t-test. n = 80 worms from 2 independent experiments. c, (top) Average ratio response of Meteor following refeeding. (bottom) Starvation and refeeding time course being that was monitored. 0 min vs. 90 min * P = 0.0246 by unpaired two-tailed t-test. n = 60 worms from 2 independent experiments.

## Discussion

In this work, we describe the development of a green fluorescent protein-based biosensor for the intracellular detection of methionine, termed Meteor. Meteor consists of a cpsfEGFP inserted into the bacterial methionine periplasmic binding protein, MetQ, and exhibits a ratiometric change in fluorescence upon binding to methionine. Previous FRET-based sensors for methionine were limited in multiplexing capacity, have low dynamic ranges, and were not used in mammalian systems^24–25^. Thus, we previously lacked the ability to measure methionine in living mammalian cells and make meaningful insights into the dynamic regulation of methionine metabolism. Meteor is the first green fluorescent protein-based biosensor to enable the study of compartmentalized uptake and synthesis of methionine in individual cells and multi-cellular organisms, resulting in the measurement of methionine dynamics across scales. Meteor demonstrates a robust selectivity for methionine and a large dynamic range, allowing for detection of methionine changes at a subcellular level and within a whole animal. Additionally, we demonstrate the utility of Meteor for multi-modal assay formats, expanding the potential uses for Meteor to study methionine dynamics in multiple contexts.

We conducted extensive characterization of Meteor, finding it to be highly specific for methionine with a K_D_ of approximately 1514 µM. Because of this lower affinity for methionine, Meteor avoids saturation in environments with abundant methionine and can report more subtle changes in methionine at biological concentrations^32–33^. This enables more dynamic tracking of methionine in cells. Additionally, our *in situ* calibration in HeLa cells revealed that Meteor responds robustly to physiological concentrations of methionine. However, while Meteor was able to capture shifts in methionine synthesis, our perturbations to the methionine cycle caused shifts that only modestly changed Meteor response, suggesting low µM concentration changes in methionine. Future efforts will involve modifying the apparent affinity of Meteor for methionine to widen the dynamic range at these smaller, physiologically relevant concentrations and improve reporting of the intracellular dynamics of methionine.

With Meteor, we can measure localized methionine dynamics in single cells, providing an alternative or tandem approach to bolster methionine metabolism studies, which have historically been limited to bulk assays. For instance, with Meteor, the dynamic, real-time changes in methionine levels can be studied with single cell resolution. Cell-to-cell heterogeneity in a cell population can be illuminated, which could be especially helpful when investigating complex environments or understanding how cell type or cell state influences methionine use. Methionine sits within the more complex methionine cycle, which is highly interconnected with other metabolic pathways^1–2^. Thus, using Meteor alongside approaches like metabolomics would enable both dynamic measurement of methionine levels in cells alongside measurements of methionine fate and changes in connected pathways^46^. While examples combining the use of biosensors with metabolomics are currently limited, we envision future integration of the two approaches would greatly enhance our understanding of metabolic dynamics.

In this study, we illuminate, for the first time, the subcellular dynamics of methionine in multiple cell types. Methionine is imported into cells using solute carriers (SLC) including SLC7A5, SLC38A1, SLC43A2, and in kidney, SLC6A18^47^. We find that exogenous methionine is found in not only the cytoplasm, cytoplasmic-facing side of the endoplasmic reticulum, and the nucleus, but also the mitochondrial matrix. Most of the methionine cycle is thought to occur in the cytoplasm, but more recently, a MAT has been found to be localized to the mitochondria for local conversion of methionine to S-adenosylmethionine^39^. As there is limited reporting on mitochondrial MATs, and how methionine would be imported to the mitochondria are unclear, future investigations into the import and use of mitochondrial methionine are warranted, which can be aided by Meteor.

The targetability of Meteor enables more precise measurements of subcellular methionine cycle dynamics. To do this, we focused on methionine dynamics in the cytoplasm and nucleus, as these are key locations for methionine cycle due to high demand for methylation reactions and maintenance of redox homeostasis. We were able to investigate the cell-to-cell variability and cell line-specific differences between HeLa, HepG2, and IHH methionine synthesis and dynamics. For example, HeLa cells were more efficient at synthesizing methionine from both hcy and MTA than HepG2 and IHH cells, with cell-to-cell variability in methionine production dynamics demonstrated across all cell populations. Within an unsynchronized cell population, the variability in response could be the result of disparities in metabolic need or cell state^48^. A potential future direction could involve using Meteor to investigate methionine metabolic demands at specific stages of the cell cycle or different cell states. Across cell types, the variability in response might be indicative of differences in how each cell type utilizes methionine and directs its metabolic flux.

Methionine restriction is reported to increase the lifespan of several model organisms, including C. elegans^12,45^. Due to the large dynamic range and good sensitivity of Meteor, we applied Meteor to sensing the dynamics of methionine in the intestine of *C. elegans*. We find that we can measure the real-time dynamics of both methionine depletion and uptake in *C. elegans*. While not explored here, these studies set the groundwork for studying more precisely the role of methionine metabolism in *C. elegans* lifespan, and we can now measure methionine dynamics during aging, which will enhance our knowledge of key biological functions and better elucidate how methionine restriction expands lifespan.

While not investigated here, the methionine cycle interconnects with both serine and cysteine metabolism. As there are recently reported GFP-based biosensors for both serine and cysteine, a red-shifted Meteor would enable multiplexed imaging of amino acid metabolism, resulting in an enhanced understanding of cell function^49–50^. Red fluorescent proteins have been used to make red-shifted calcium, cyclic AMP, and lactate reporters, and could be incorporated into Meteor^51–53^. Similarly, the self-labeling protein HaloTag has been used to make far-red calcium and kinase biosensors and could be adapted into the Meteor design for a far-red biosensor for multiplexed, compartmentalized sensing of metabolic dynamics^54–55^.

Overall, we have developed Meteor, a sensitive, single-fluorophore biosensor for methionine, which can be used to measure methionine dynamics with high spatial and temporal resolution across scales. With Meteor, measurement of methionine dynamics in single mammalian cells and multi-cellular organisms is now possible, greatly enhancing our set of tools to measure metabolism across scales in real time.

## Methods

### Chemicals

L-methionine (Sigma-Aldrich, Cat# M-9625), L-homocysteine (AA Blocks, Cat# AA0037CV), 5’-methylthioadenosine (MedChem Express, Cat# HY-16938-100mg), and sodium methotrexate (Thermo Scientific, Cat# J66364-MD) were dissolved in PBS. L-alanine (Thermo Scientific, Cat# AAA1580422), L-arginine (Thermo Fisher Scientific, Cat# A15738-14), L-asparagine (Thermo Scientific Cat# AAB2147336), L-aspartic acid (Thermo Scientific, Cat# AAA1352030), L-cysteine (Millipore Sigma, Cat# 2430-M), L-glutamic acid (Thermo Scientific, Cat #AC156212500), L-glutamine (Thermo Scientific, Cat# AAA1420130), glycine (Sigma-Aldrich, Cat# 50046), L-histidine (Sigma-Aldrich, Cat# H-8125), L-isoleucine (Fisher Scientific, Cat# BP384), L-leucine (Fisher Scientific, Cat# BP385), L-lysine (Millipore Sigma, Cat# 4400-M), L-phenylalanine (Sigma-Aldrich, Cat# P-2126), L-proline (Sigma-Alrich, Cat# P-0380), L-serine (Sigma-Aldrich, Cat# S4500), L-threonine (Sigma-Aldrich, Cat# T-8625), L-tryptophan (Sigma-Aldrich, Cat# T0254), L-tyrosine (Sigma-Aldrich, Cat# T-3754), and L-valine (Fisher Scientific, Cat# BP397) were dissolved in PBS.

NmMetQ WT was a gift from Douglas Rees (Addgene plasmid # 129647; http://n2t.net/addgene:129647; RRID:Addgene_129647)s.

### Plasmids

Initial insertion designs were cloned by Gibson Assembly (Gibson Assembly HiFi Master Mix, Fisher Scientific, Cat# A46628) to insert cpsfEGFP with SAG-GT linkers into a pRSET vector that contained MetQ^56^. cpsfEGFP was amplified from iGlo.

To create linker libraries, Golden Gate Assembly was used^57^. Primers contained degenerate NNK codons at mutated positions. PCR was performed with MetQ51 as template and subsequently digested with DpnI (Thermo Scientific, Cat# FD1703). PCR cleanup was performed (Thermo Scientific, Cat# K310001), and the resulting fragments were digested with BsmBI and then ligated. The product was transformed into DH5*α* cells (Thermo Scientific, Cat# 18265017). The transformation was diluted in LB containing 100 μg/mL ampicillin for overnight growth, and a small aliquot was plated on LB plates containing 100 μg/mL ampicillin to monitor the number of colony forming units (cfu) produced. The overnight culture was miniprepped (Qiagen, Cat# 27106) to get a library of linker variants.

To make targeted Meteor, localization sequences were appended to either the N-terminus or C-terminus of Meteor using Gibson Assembly. Localization sequences can be found in Supplementary Table 1. MeteorITE libraries were created using degenerate NNK codons at mutated amino acid residues and Golden Gate Assembly. To make Meteor-mScarlet, mScarlet was amplified from Malibu-mScarlet and appended to the C-terminus of Meteor using Gibson Assembly^58^.

### Meteor Sequence

Linkers are underlined in the following sequence: MKEIVFGTTVSCWNVYIMADKQKNGIKANFKIRHNVEDGSVQLADHYQQNTPIGDGPVLLPDNHYLSTQSVLSKDPNEKRDHMVLLEFVTAAGITLGMDELYKGGTGGSMSKGEELFTGVVPILVELDGDVNGHKFSVRGEGEGDATNGKLTLKFICTTGKLPVPWPTLVTTLTYGVQCFSRYPDHMKQHDFFKSAMPEGYVQERTISFKDDGTYKTRAEVKFEGDTLVNRIELKGIDFKEDGNILGHKLEYNLQGDFGDMVKEQIQAELEKKGYTVKLVEFTDYVRPNLALAEGELDINVFQHKPYLDDFKKEHNLDITEVFQVPTAPLGLYPGKLKSLEEVKDGSTVSAPNDPSNFARVLVMLDELGWIKLKDGINPLTASKADIAENLKNIKIVELEAAQLPRSRADVDFAVVNGNYAISSGMKLTEALFQEPSFAYVNWSAVKTADKDSQWLKDVTEAYNSDAFKAYAHKRFEGYKSPAAWNEGAAK*

### Lysate Screening

To screen the initial insertion site variants, plasmids were transformed into BL21 *E. coli* and plated on LB-ampicillin plates. The following day, colonies were inducted in 10 mL of ZYM-5052 autoinduction media with ampicillin^59^. Cultures were grown at 37°C and 250 rpm shaking for 6 h, followed by an incubation at 20°C and 250 rpm shaking for 22 h. Cultures were pelleted by centrifugation and then resuspended in PBS at 20°C and 250 rpm shaking for 1 h to reduce residual exogenous methionine levels. Cultures were pelleted again and lysed in B-PER lysis buffer (Thermo Scientific Cat# P178248) containing protease inhibitor (Thermo Scientific Cat# A32963). The lysate was clarified by centrifugation, and the supernatant was used for fluorescence reading by plate reader (Molecular Devices SpectraMax iD5) in a 96-well plate (Genesee Scientific, Cat# 33-755). The plate reader was equipped with SoftMax Pro 7.1 data acquisition software (Molecular Devices). Fluorescence readings were measured with 405 nm and 480 nm excitation and with 520 nm emission before and after adding 100 µM methionine to the lysate. ΔF/F was calculated by taking the difference between the final and initial fluorescence values and dividing by the initial fluorescence value. ΔR/R was calculated in a similar manner but using a ratio instead of raw fluorescence. This ratio was calculated by dividing the 480 nm excitation-emission by 405 excitation-emission.

For linker screenings, linker libraries were transformed in BL21 cells and plated on LB-ampicillin plates. For each round of linker screening, 192 colonies were selected and inoculated in 1 mL of ZYM-5052 auto induction media in deep well culture plates (Genesee Scientific, Cat# 27-413), covered with a sterile sealing film (Genesee Scientific, Cat # 12-631), and cultured as previously described. Once cultures were pelleted and lysed, the resulting supernatant was transferred to a 96-well assay plate. Fluorescent measurements were taken before and after addition of 100 µM methionine.

### Protein Purification

A 6xHis tag and TEV cleavage site were added to the N-terminus of Meteor and MeteorITE in a pRSET vector. BL21 *E. coli* transformed with these plasmids were cultured in OM-I media and induced with 100 µM IPTG (Goldbio, Cat# I2481C50)^60^. Cells were harvested by centrifugation and then mechanically lysed using an EmulsiFlex-C3 (Avestin) in lysis buffer (50 mM Tris, 150 mM NaCl, 20 mM imidazole, 5% glycerol, pH 8.0). The lysate was clarified by centrifugation, and the supernatant was incubated with Ni-NTA resin (Thermo Scientific, Cat# 25215). The resin washed with lysis buffer and incubated with elution buffer (50 mM Tris, 150 mM NaCl, 500 mM imidazole, 5% glycerol, pH 8.0) before the eluate was collected. Eluate was concentrated in 10 kDa MWCO concentrators (Thermo Scientific Cat# 88528). 0.3 mg of MBP-super TEV protease was added to the concentrated eluate to remove the 6xHis tag, and the samples were dialyzed with 10 kDa MWCO membrane (Thermo Scientific Cat# PI88243) in protein storage buffer (50 mM Tris, 150 mM NaCl, 5% glycerol, pH 8.0). The digested and dialyzed eluate was incubated again with Ni-NTA resin, washed with protein storage buffer, and the final flow through and all fractions were assessed via SDS-PAGE (ThermoFisher, Cat# NP0322BOX). Purified protein was snap frozen in liquid N_2_ and stored in -80 °C.

### In Vitro Meteor Assays

*In vitro* characterization of Meteor and MeteorITE was performed in 96-well opaque, black assay plates (Genesee Scientific, Cat# 33-755). All *in vitro* assays used 5 uM protein in protein assay buffer (50 mM Tris, 150 mM NaCl, 5% glycerol, variable pH). For dose-response and specificity assays, assay buffer with pH 7.2 was used. For pH sensitivity assays, assay buffered were adjusted to pH 6.5, 7.0, 7.2, 7.5, 7.7 and 8.0. *K*_D_ values were calculated by nonlinear fit using equation 1 in GraphPad Prism 10.

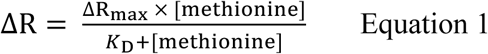

All assays were completed in a plate reader with fluorescence read from the top using monochromators with 405 nm and 480 nm excitation wavelengths and 520 nm emission. For excitation spectra, emission was detected at 550 nm. For emission spectra, excitation was conducted at 405 nm and 480 nm. Detection used an ultracooled photomultiplier tube.

Stopped flow kinetics experiments were performed using an Applied Photometrics SX20. Meteor and methionine in pH 7.2 protein assay buffer were mixed to a final concentration of 300 nM Meteor and 31.6 mM methionine. Experiments were conducted at 37 °C. An excitation wavelength of 480 nm was used and a bandpass of 6.98 nm was set using a monochromator (Applied Photophysics). Emission was filtered through a 515 nm longpass filter (CVI Laser). Fluorescence was measured every 360 ms for 360 seconds with detector voltage set to 300 V. Data was collected using the Pro-Data SX data acquisition software (Applied Photophysics). K_obs_ values were determined using one-phase association in GraphPad Prism 10.

### Cell Culture and Transfection

HeLa (ATCC Cat# CRM-CCL-2), U2OS (ATCC Cat# HTB-96), HepG2 (ATCC Cat# HB-8065), and IHH (provided by Thomas Vallim) were cultured at 37 °C with 5% CO_2_ in Dulbecco’s modified Eagle medium (DMEM, Thermo Scientific Cat# 10569010) containing 4.5 g/L glucose, glutaMAX supplement, sodium pyruvate, 10% fetal bovine serum (Avantor 76419-584), and 100 U/mL penicillin-streptomycin (Fisher Scientific Cat# 15-140-122). Routine mycoplasma checks were performed using NucBlue staining (Fisher Scientific Cat# R37605) and PCR testing with a PCR mycoplasma detection kit (Fisher Scientific Cat# AAJ66117AMJ).

HeLa and U2OS cells were seeded at 150k-200k per dish one day prior to transfection or 75k-100k two days prior. HepG2 and IHH cells were seeded at 300k per dish one day prior to transfection or 150k two days prior. Cells were seeded in 35 mm glass bottom dishes (Cellvis, Cat# d35-14-1.5-n). Transfections for HeLa and U2OS were conducted one day prior to imaging using 500 ng plasmid DNA, Opti-MEM (Gibco Cat#31985070), and FuGENE HD (Promega Cat#HD-1000) following manufacturer’s protocol. Transfections for HepG2 and IHH were conducted one day prior to imaging using 600 ng plasmid DNA, Opti-MEM, and FuGENE 4K (Promega Cat#E2311).

### Fluorescence Imaging and Image Analysis

Fluorescence microscopy experiments were performed using a Nikon ECLIPSE Ti2 epifluorescence microscope with a CF160 Plan Fluor 40x Oil Immersion Objective Lens (N.A. 1.3, W.D. 0.2 mm, F.O.V. 25 mm; Nikon), a Spectra III UV, V, B, C, T, Y, R, nIR light engine with 380/20, 475/28, and 575/25 LEDs or Spectra III, Custom Spectra III filter sets (440/510/575 and 390/475/555/635/747) stationed in a Ti cube polychroic, a Kinetix 22 back-illuminated sCMOS camera (Photometrics), and a stage-top incubator set to 37 °C (Tokai Hit). The microscope is controlled with NIS-Elements software (Nikon). Exposure times ranged from 100 to 200 ms for all Meteor and MeteorITE constructs.

Cells were washed three times with 2mL HBSS (ThermoFisher Scientific Cat# 14065056 supplemented with 2 g/L glucose, 20 mM HEPES, pH 7.4) and incubated at 37 °C with HBSS for 30 minutes prior to imaging. For experiments involving methionine starvation, DMEM was exchanged for methionine-depleted DMEM (ThermoFisher Scientific Cat# 21013024) supplemented with cysteine (Sigma-Aldrich Cat# C7602-25G) and dFBS (ThermoFisher Scientific Cat#26400044), and cells were incubated at 37 °C for two hours prior to imaging. Images were collected using a 40x objective and were taken every 30 seconds for 30 minutes. Additions were made after four minutes of baseline imaging. Either 150 µM homocysteine (AA Blocks, Cat# AA0037CV) or 30 µM 5’-methylthioadenosine (MedChem Express, Cat# HY-16938-100mg) were added to cells. For methotrexate experiments, cells were incubated in methionine-depleted DMEM supplemented with cysteine and dFBS with 100 µM methotrexate (Thermo Scientific, Cat# J66364-MD) at 37 °C for two hours prior to imaging.

Image analysis was performed as previously described using MATLAB R2024a (MathWorks)^58^. Cell or dishes that were not efficiently transfected, dying, or under or over exposed were excluded from this analysis.

### Confocal Imaging

HeLa cells expressing localized Meteor variants were imaged using confocal microscopy on a Leica Stellaris 5 DMi8 Confocal Microscope with DMOD WLL (440 nm – 790 nm), HC PL APO 63x/1.40 oil immersion CS2 lens, and a Power HyD S detector. LasX software (Leica) controlled the microscope, and 0.5-8% laser intensities were used to image the cells. Cells expressing cytoplasmic- and nuclear-localized Meteor were washed with HBSS and stained with NucBlue (Fisher Scientific, Cat# R37605) for 15 minutes prior to imaging. Cells expressing outer endoplasmic reticulum and mitochondrial matrix Meteor were washed with HBSS and stained with 25 nM ERTracker Deep Red (Fisher Scientific, Cat# E34250) or MitoTracker Deep Red (Thermo Scientific, Cat# M22426), respectively.

### Plate Reader-Based Mammalian Assay

1.5x10^6^ HeLa cells were seeded in 10 cm cell culture dishes (Genesee Scientific Cat# 25-202). After 24 hours, cells were transfected with FuGENE HD (Promega Cat# HD-1000) and 2.5 μg of either Meteor-mScarlet or MeteorITE-mScarlet plasmid according to the manufacturer’s protocol. After an additional 24 hours, cells were dissociated with TrypLE Express (ThermoFisher Scientific Cat# 12604013). Trypsin was inactivated with Dulbecco’s modified Eagle medium (DMEM, Thermo Scientific Cat# 10569010) containing 4.5 g/L glucose, glutaMAX supplement, sodium pyruvate, 10% fetal bovine serum (Avantor 76419-584), and 100 U/mL penicillin-streptomycin (Fisher Scientific Cat# 15-140-122). The cell suspension was centrifuged at 2000 g for 3 minutes. The supernatant was aspirated, and the cell pellet was resuspended in DMEM. 30,000 Meteor-mScarlet or MeteorITE-mScarlet cells were seeded in each well in 96-well plates with clear wells (Genesee Scientific Cat# 22-721) and incubated at 37 °C overnight.

Once adherent, cells were washed three times with 2mL HBSS (ThermoFisher Scientific Cat# 14065056 supplemented with 2 g/L glucose, 20 mM HEPES, pH 7.4) and incubated at 37 °C with HBSS for 30 minutes prior to the plate reader assay. The plate reader was equipped with SoftMax Pro 7.1 data acquisition software (Molecular Devices). Fluorescence readings were measured at 405 nm and 480 nm excitation with 520 nm emission before and after adding no methionine, 10 µM methionine, or 100 µM methionine to each well. The excitation-emission ratio was calculated by equation 2.

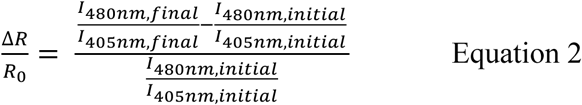

### Flow Cytometry

1.5x10^6^ HeLa cells were seeded in 10 cm cell culture dishes (Genesee Scientific Cat# 25-202). After 24 hours, cells were transfected with FuGENE HD (Promega Cat# HD-1000) and 2.5 μg of either Meteor-mScarlet or MeteorITE-mScarlet plasmid according to the manufacturer’s protocol. After an additional 24 hours, cells were washed with HBSS (ThermoFisher Scientific Cat# 14065056 supplemented with 2 g/L glucose, 20 mM HEPES, pH 7.4) and dissociated with TrypLE Express (ThermoFisher Scientific Cat# 12604013). Trypsin was inactivated with Dulbecco’s modified Eagle medium (DMEM, Thermo Scientific Cat# 10569010) containing 4.5 g/L glucose, glutaMAX supplement, sodium pyruvate, 10% fetal bovine serum (Avantor 76419-584), and 100 U/mL penicillin-streptomycin (Fisher Scientific Cat# 15-140-122). The cell suspension was centrifuged at 2000 g for 3 minutes. The supernatant was aspirated, and the cell pellet was resuspended in HBSS and divided into samples containing no methionine, 30 μM methionine, or 300 μM methionine.

Cell suspensions were passed through a cell strainer (Fisher Scientific Cat# 08-771-23) prior to loading onto a flow cytometer (Bio-Rad S3 Cell Sorter, Cat# 145-1002). The instrument was set to maintain a temperature of 37 °C, collect 50,000 events per run, and use both 488 nm and 561 nm lasers. cpsfEGFP fluorescence was measured with 525/30 emission (FL1), while mScarlet fluorescence was measured using 586/25 emission (FL2). FlowJo V10 was used for data analysis. A single population of cells was gated with FSC (forward scatter) and SSC (side scatter). This population was then gated with FL2 such that there were minimal overexposed events in FL1. Plots showing gated FL1 and FL2 fluorescence values were plotted in FlowJo v10. Fluorescence values were exported and the average FL1/FL2 ratio (cpsfEGFP/mscarlet) from each biological replicate was plotted in GraphPad Prism 10.

### Immunostaining

Cells were seeded and grown to 70-80% confluency in 6-well glass bottom dishes (Cellvis Cat #P06-1.5H-N) prior to treatment with small molecules. Following treatment, cells were washed with cold 1x PBS (Thermo Fisher Cat #10010023) three times for five minutes and then fixed with 4% (v/v) paraformaldehyde (Fisher Scientific Cat #50-980-487) for 20 min at room temperature. Post-fixation, cells were washed three times for 5 minutes each with 1x PBS and then permeabilized in 1x PBS with 0.1% Triton X-100 (Thermo Fisher Cat #A16046.AE) and 5% BSA (Tocris Biosciences, Cat #5217) per well. After blocking, methionine synthase (MTR) rabbit polyclonal antibody (Abcam Cat #AB66039) was added to the wells at a 1:500 dilution and rocked overnight at 4°C. The next day, cells were washed three times for five minutes each with 1x PBS and then incubated and rocked with HRP-goat anti-rabbit IgG (H+L) secondary antibody conjugated to Alexa Fluor Plus 488 (1:200; Thermo Scientific Cat #A-32731), protected from light, for 1 hour at room temperature. Post-incubation, the cells were washed three times for five minutes each with 1x PBS, and NucBlue (Thermo Fisher Cat #R37605) was added during the second wash. The dishes were imaged one day post using confocal microscopy.

Images were quantified using a CellProfiler pipeline. The pipeline first detects individual nuclei stained with NucBlue from the images of nuclei alone using global Otsu thresholding and shape-based separation of touching nuclei. It then rescales the image containing the immunostained protein of interest and uses this rescaled image, along with the detected nuclei as seeds, to propagate outward and define whole-cell boundaries. Any cells touching the image border are discarded along with their corresponding nuclei. The cytoplasmic compartment is then defined as the difference between the whole-cell area and the nuclear area, yielding a ring-shaped region for each cell. Mean fluorescence intensity of the cytoplasm is measured separately within the nuclear and cytoplasmic regions of each cell, and these two values are divided to produce the N/C ratio.

### C. elegans

Meteor was expressed in the *C. elegans* intestine using an intestine-specific *vha-6* promoter^61^. An expression vector based on pCFJ356 plasmid containing the *vha-6* promoter, mRuby, *unc-54* 3’UTR, and *cb-unc-119(+)* was PCR amplified to have the *vha-6* promoter and mRuby at the ends. A fragment containing a codon optimized Meteor sequence with two inserted intron sequences and 40 bases of overlapping sequences with the vha-6 promoter and mRuby, was ordered from Twist Bioscience. The *C. elegans* expression vector for Meteor was then assembled by Gibson cloning. Microinjections were performed on *unc-119(ed3)* mutant animals with a mixture of 25 ng/μL of the Meteor expression plasmid and 75 ng/μL of an empty expression vector as filler DNA. Meteor expression strains were maintained by selecting for wild-type animals and maintaining at 15°C on OP50 *E. coli* strain grown on NGM plates (Bacto-Agar (Difco) 2% w/v, Bacto Peptone 0.25% w/v, NaCl_2_ 0.3% w/v, 1 mM CaCl_2_, 5 µg/ml cholesterol, 0.625 mM KPO_4_ pH 6.0, 1 mM MgSO_4_).

For experiments, animals were synchronized using a standard bleaching protocol. Worms were collected using M9 solution (22 mM KH_2_PO_4_ monobasic, 42.3 mM NaHPO_4_, 85.6 mM NaCl, 1 mM MgSO_4_) into a 15 mL conical tube. M9 was then replaced with a bleaching solution (1.8% sodium hypochlorite, 0.375 M NaOH in M9) at a 20:1 bleach:worm volume ratio and vigorously shaken until all worms were dissolved and only intact eggs remain. Eggs were then washed four times in M9 solution by centrifugation at 1,100 x g for 1 min. After the final wash, animals were L1 arrested by incubating overnight in M9 solution on a rotator at 20°C. L1 animals were grown on OP50 bacteria on NGM plates until day 1 of adulthood (3 days at 20°C). Animals were washed 3x with M9 and starved in M9 solution at 20°C on a tube rotator for the designated times and refed by returning animals onto OP50 bacteria on NGM plates.

For imaging, animals were manually picked onto a drop of M9 containing 100 mM sodium azide on an NGM plate without bacteria. Once the M9 dried, the immobilized animals were arranged in the same orientation and imaged on an M205 Leica stereomicroscope with a Leica LED3 light source, standard GFP filter, and DsRed filter. Starved animals were first transferred to an NGM plate without bacteria to be picked for imaging. Collected images were analyzed in FIJI, where an ROI was drawn around each worm using the DIC image, and mean intensities of the ROI from both the Meteor image and RFP image quantified. A background ROI with no *C. elegans* was collected. The background-corrected ratio of Meteor 480 nm ex-em to RFP was then calculated.

### Statistics and Reproducibility

Figure preparation and statistical analysis were performed using GraphPad Prism 10, except for Fig 3c-d which were prepared in FlowJo v10. The statistical tests used and P values are reported in all figure legends where applicable. The number of cells analyzed (n cells), and number of independent experiments are reported in all figure legends. For imaging experiments, statistics were performed using the means of all single cell measurements from independent experiments. All time courses shown are the mean plus the standard deviation for all cells unless otherwise noted. All dot plots shown depict the mean ± standard deviation. When applicable, data were plotted using the SuperPlots method^62^. All time course data was baseline corrected by fitting the baseline preaddition to a linear model and subtracting predicted values from those observed.

## Acknowledgements

We thank the following: Martin Phillips for his assistance with stopped flow, Mark Arbing and UCLA-DOE Protein Expression Technology Center for access to a Bio-Rad S3 cell sorter which is supported by Department of Energy Grant DE-FC02-02ER63421, Jack Scully for his assistance with protein purification and stopped flow, Peter DePaola for his assistance with protein purification and helpful discussion, Thomas Vallim for providing IHH cells, Michael Lawson for material support, Tara TeSlaa and all members of the Schmitt Lab for their helpful discussion and critique.

This work was supported by the National Institutes of Health (DP2GM154012-01 to D.L.S R01AG079806 to R.H-S and R01AG079806-02S1 to G.G.), the Chan-Zuckerberg Initiative (MET-000000000151 to D.L.S.), UCLA Jonsson Comprehensive Cancer Center Graduate Research Award (to K.A.C.), UCLA Eli and Edythe Broad Center of Regenerative Medicine and Stem Cell Research Transformative Technology Development Award, including support from Carol Doumani (to D.L.S.).

## Author Contributions

D.L.S. and K.A.C. conceptualized and planned the project. K.A.C developed Meteor. K.A.C., A.J., N.G., and M.C. performed and analyzed all time course imaging. K.A.C and M.C. performed flow cytometry. A.J. performed immunostaining. G.G., A.A., and R.H-S. prepared and performed all experiments with *C. elegans*. D.L.S. and R.H-S supervised all work. K.A.C. and D.L.S. analyzed all data. K.A.C. prepared the figures. K.A.C. and D.L.S. wrote the manuscript. All authors discussed results and approved the final version of the manuscript.

## Resource Availability

All plasmids encoding Meteor and MeteorITE will be deposited with Addgene. Meteor-expressing *C. elegans* will be deposited with the Caenorhabditis Genetics Center (CGC).

