## Supplemental Information for "Revealing the spatiotemporal dynamics of methionine metabolism with a genetically encoded single-fluorophore biosensor"

### Supplemental Figures and Tables

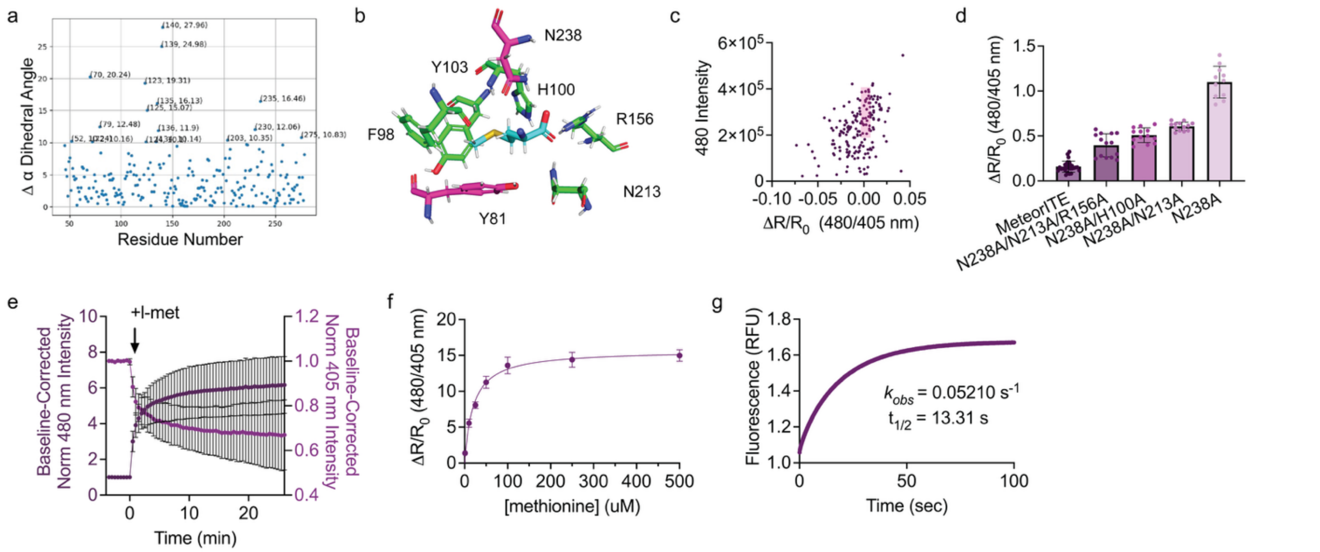


**Supplementary Figure 1. Development and characterization of Meteor and MeteorITE**

a, Calculated dihedral angle change of the α carbon experienced at each amino residue when MetQ binds to methionine. The apo (PDB ID 6CVA) and holo (PDB ID 6OJA) conformations of MetQ were compared, and residues experiencing an angle change greater than 10° are annotated.

b, Representative binding pocket for methionine-bound MetQ (PDB ID 6OJA) highlighting key amino acid residues that interact with methionine. Methionine is shown in cyan, and the sites ultimately mutated to eliminate binding affinity in MeteorITE are shown in pink (Y81 and N238).

c, Screen of a variant library created by site saturation mutagenesis at residues Y81 and N238. The maximum ratio change ($\Delta R/R_0$) of variants was plotted versus 480 nm intensity when screened in clarified bacterial lysate with 1 mM methionine to identify variants that are both bright but do not respond to methionine. The pink box highlights variants selected for further screening towards a Meteor variant with no binding affinity for methionine.

d, Maximum ratio change of MeteorITE and select MeteorITE candidates in response to 30 μM methionine. MeteorITE n = 27 cells from 2 independent experiments. N238A/N213A/R156A n = 14 cells from 2 independent experiments. N238A/H100A n = 13 cells from 2 independent experiments. N238A/N213A n = 13 cells from 2 independent experiments. N238A n = 11 cells from 2 independent experiments.

e, Fluorescence changes at 480 nm (dark purple) and 400 nm (magenta) in response to 30 μM methionine over single-cell time course imaging in HeLa cells expressing Meteor.

f, Fluorescence vs. time of 500 nM Meteor mixed with the 31.6 mM methionine measured by stopped flow. The graph is indicative of the average of 4 technical replicates. $K_{obs}$ was calculated using a one-phase association curve.

g, The maximum ratio change Meteor in response to various concentrations of methionine (from 1 µM to 500 µM) in HeLa cells permeabilized with 0.0005% digitonin. The graph is indicative of the average of 10 to 14 cells per concentration.

For all graphs, mean ± standard deviation is shown.

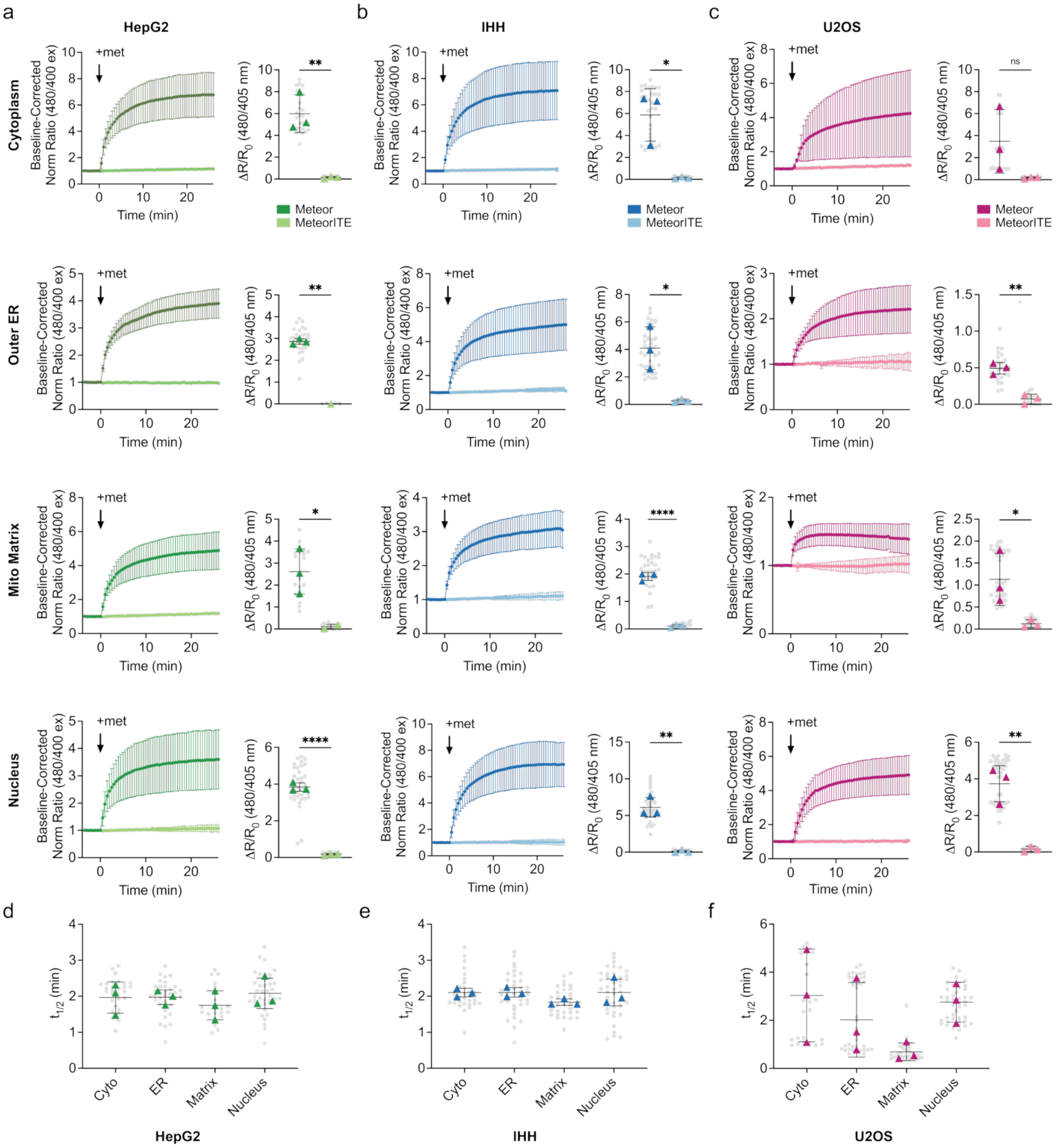


**Supplementary Figure 2. Meteor measures subcellular methionine uptake in HepG2, IHH, and U2OS cell lines**

a, (left) Normalized response of localized Meteor (dark green) and MeteorITE (light green) to 30 μM methionine addition at (from top to bottom) the cytoplasm, outer endoplasmic reticulum, mitochondrial matrix, and nucleus in HepG2 cells as well as the (right) average responses of Meteor and MeteorITE at each location. Cyto ** $P = 0.0044$; ER ** $P = 0.0026$; matrix * $P = 0.0473$; nucleus **** $P < 0.0001$ by unpaired two-tailed t-test. Cyto-Meteor n = 27 cells from 3 independent experiments. Cyto-MeteorITE n = 24 from 3 independent experiments. ER-Meteor n = 36 cells from 3 independent experiments. ER-MeteorITE n = 13 from 3 independent experiments. Matrix-Meteor n = 21 cells from 3 independent experiments. Matrix-MeteorITE n = 21 from 3 independent experiments. NLS-Meteor n = 36 cells from 3 independent experiments. NLS-MeteorITE n = 42 from 3 independent experiments.

b, (left) Normalized response of localized Meteor (dark blue) and MeteorITE (light blue) to 30 μM methionine addition at (from top to bottom) the cytoplasm, outer endoplasmic reticulum, mitochondrial matrix, and nucleus in IHH cells as well as the (right) average responses of Meteor and MeteorITE at each location. Cyto * P = 0.0142; ER * P = 0.0131; matrix **** P < 0.0001; nucleus ** P = 0.0014 by unpaired two-tailed t-test. Cyto-Meteor n = 32 cells from 3 independent experiments. Cyto-MeteorITE n = 26 from 3 independent experiments. ER-Meteor n = 39 cells from 3 independent experiments. ER-MeteorITE n = 24 from 3 independent experiments. Matrix-Meteor n = 33 cells from 3 independent experiments. Matrix-MeteorITE n = 20 from 3 independent experiments. NLS-Meteor n = 43 cells from 3 independent experiments. NLS-MeteorITE n = 37 from 3 independent experiments.

c, (left) Normalized response of localized Meteor (dark pink) and MeteorITE (light pink) to 30 μM methionine addition at (from top to bottom) the cytoplasm, outer endoplasmic reticulum, mitochondrial matrix, and nucleus in U2OS cells as well as the (right) average responses of Meteor and MeteorITE at each location. Cyto ns P = 0.1201; ER * P = 0.0432; matrix ** P = 0.0021; nucleus ** P = 0.0034 by unpaired two-tailed t-test. Cyto-Meteor n = 25 cells from 3 independent experiments. Cyto-MeteorITE n = 30 from 3 independent experiments. ER-Meteor n = 38 cells from 3 independent experiments. ER-MeteorITE n = 31 from 3 independent experiments. Matrix-Meteor n = 28 cells from 3 independent experiments. Matrix-MeteorITE n = 20 from 3 independent experiments. NLS-Meteor n = 43 cells from 3 independent experiments. NLS-MeteorITE n = 33 from 3 independent experiments.

d, Time-to-half-maximum (t1/2) of subcellular targeted Meteor response to 30 μM methionine in live HepG2 cells from experiments shown in Fig S2a. n.s. P = 0.7528 ordinary one-way ANOVA; pairwise comparisons are all n.s. P > 0.05 by Tukey's multiple comparisons test.

e, Time-to-half-maximum (t1/2) of subcellular targeted Meteor response to 30 μM methionine in live IHH cells from experiments shown in Fig S2b. n.s. P = 0.1078 ordinary one-way ANOVA; pairwise comparisons are all n.s. P > 0.05 by Tukey's multiple comparisons test.

f, Time-to-half-maximum (t1/2) of subcellular targeted Meteor response to 30 μM methionine in live U2OS cells from experiments shown in Fig S2c. n.s. P = 0.2094 ordinary one-way ANOVA; pairwise comparisons are all n.s. P > 0.05 by Tukey's multiple comparisons test.

For all graphs, individual cells are shown in small, gray circles, and the average of each biological replicate is shown in a colored triangle.

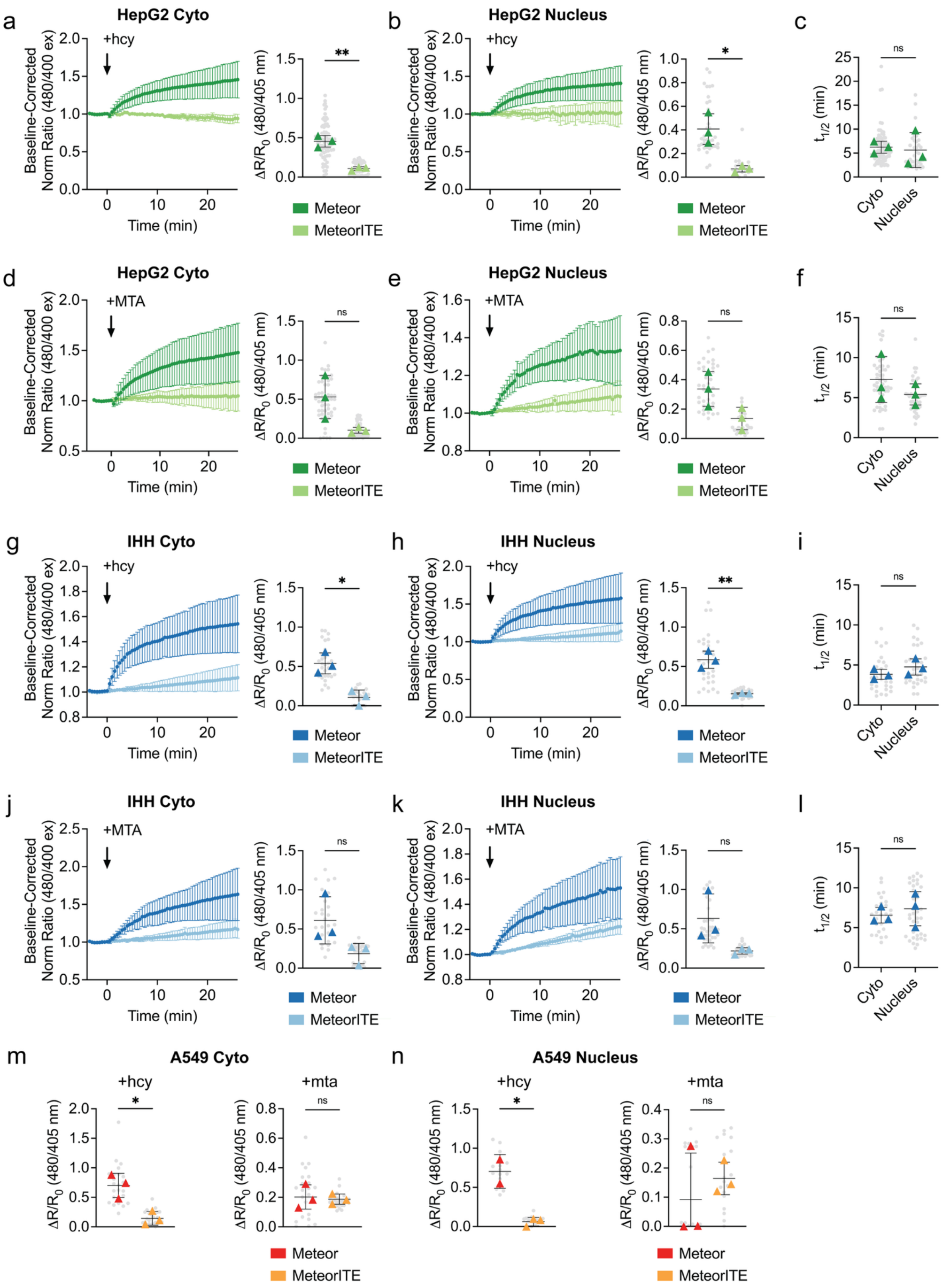

a
HepG2 Cyto
+hcy
Baseline-Corrected Norm Ratio (480/400 ex)
Time (min)
ΔR/R0 (480/405 nm)
**
Meteor
MeteorITE
b
HepG2 Nucleus
+hcy
*
c
ns
t1/2 (min)
Cyto
Nucleus
d
HepG2 Cyto
+MTA
e
HepG2 Nucleus
+MTA
f
g
IHH Cyto
+hcy
h
IHH Nucleus
i
j
IHH Cyto
+MTA
k
IHH Nucleus
l
m
A549 Cyto
+hcy
+mta
n
A549 Nucleus

**Supplementary Figure 3 – Meteor measures subcellular methionine synthesis in HepG2, IHH, and A549 cell lines**

a-b, (left) Normalized response of localized Meteor (dark green) and MeteorITE (light green) to 150 µM homocysteine addition at either the cytoplasm or nucleus in HepG2 cells as well as the (right) average responses of Meteor and MeteorITE at either location. Cyto ** P = 0.0014; Nucleus * P = 0.0118 by unpaired two-tailed t-test. Cyto-Meteor n = 80 cells from 3 independent experiments. Cyto-MeteorITE n = 55 from 3 independent experiments. NLS-Meteor n = 37 cells from 3 independent experiments. NLS-MeteorITE n = 20 from 3 independent experiments.

c, Time-to-half-maximum (t1/2) of subcellular targeted Meteor response to 150 µM homocysteine in live HepG2 cells from experiments shown in Fig S3a-b. n.s. P = 0.7991 by unpaired two-tailed t-test.

d-e, (left) Normalized response of localized Meteor (dark green) and MeteorITE (light green) to 30 µM MTA addition at either the cytoplasm or nucleus in HepG2 cells as well as the (right) average responses of Meteor and MeteorITE at either location. Cyto n.s. P = 0.0589; Nucleus n.s. P = 0.0678 by unpaired two-tailed t-test. Cyto-Meteor n = 54 cells from 3 independent experiments. Cyto-MeteorITE n = 57 from 3 independent experiments. NLS-Meteor n = 39 cells from 3 independent experiments. NLS-MeteorITE n = 26 from 3 independent experiments.

f, Time-to-half-maximum (t1/2) of subcellular targeted Meteor response to 30 µM MTA in live HepG2 cells from experiments shown in Fig S3d-e. n.s. P = 0.3655 by unpaired two-tailed t-test.

g-h, (left) Normalized response of localized Meteor (dark blue) and MeteorITE (light blue) to 150 µM homocysteine addition at either the cytoplasm or nucleus in IHH cells as well as the (right) average responses of Meteor and MeteorITE at either location. Cyto * P = 0.0101; Nucleus ** P = 0.0025 by unpaired two-tailed t-test. Cyto-Meteor n = 30 cells from 3 independent experiments. Cyto-MeteorITE n = 27 from 3 independent experiments. NLS-Meteor n = 31 cells from 3 independent experiments. NLS-MeteorITE n = 28 from 3 independent experiments.

i, Time-to-half-maximum (t1/2) of subcellular targeted Meteor response to 150 µM homocysteine in live IHH cells from experiments shown in Fig S3g-h. n.s. P = 0.2529 by unpaired two-tailed t-test.

j-k, (left) Normalized response of localized Meteor (dark blue) and MeteorITE (light blue) to 30 µM MTA addition at either the cytoplasm or nucleus in IHH cells as well as the (right) average responses of Meteor and MeteorITE at either location. Cyto n.s. P = 0.0884; Nucleus n.s. P = 0.0847 by unpaired two-tailed t-test. Cyto-Meteor n = 26 cells from 3 independent experiments. Cyto-MeteorITE n = 31 from 3 independent experiments. NLS-Meteor n = 38 cells from 3 independent experiments. NLS-MeteorITE n = 31 from 3 independent experiments.

l, Time-to-half-maximum (t1/2) of subcellular targeted Meteor response to 30 μM MTA in live IHH cells from experiments shown in Fig S3j-k. n.s. P = 0.5790 by unpaired two-tailed t-test.

m, Average responses of Meteor and MeteorITE at the cytoplasm to either (left) homocysteine or (right) MTA. Hcy * P = 0.0147; MTA n.s. P = 0.7922 by unpaired two-tailed t-test. Meteor Hcy n = 24 cells from 3 independent experiments. MeteorITE Hcy n = 25 from 3 independent experiments. Meteor MTA n = 22 cells from 3 independent experiments. MeteorITE MTA n = 25 from 3 independent experiments.

n, Average responses of Meteor and MeteorITE at the nucleus to either (left) homocysteine or (right) MTA. Hcy * P = 0.0129; MTA n.s. P = 0.5007 by unpaired two-tailed t-test. Meteor Hcy n = 16 cells from 2 independent experiments. MeteorITE Hcy n = 20 from 3 independent experiments. Meteor MTA n = 24 cells from 3 independent experiments. MeteorITE MTA n = 19 from 3 independent experiments.

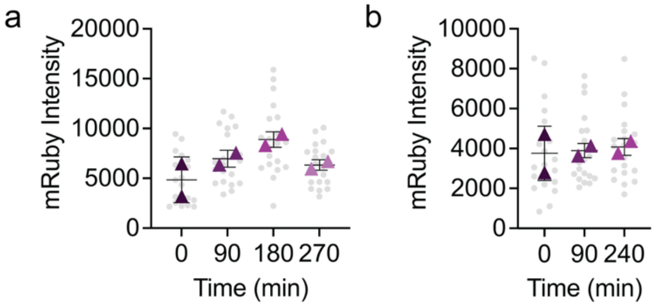


**Supplementary Figure 4 – Meteor measures *in vivo* methionine dynamics in a *C. elegans* model**

a, Background-corrected RFP intensity in *C. elegans* from Fig 6b during the starvation period. n.s. P = 0.1426 by ordinary one-way ANOVA; pairwise comparisons are all n.s. P > 0.05 by Tukey's multiple comparisons test.

b, Background-corrected RFP intensity in *C. elegans* from Fig 6c during the refeeding period. n.s. P = 0.9340 by ordinary one-way ANOVA; pairwise comparisons are all n.s. P > 0.05 by Tukey's multiple comparisons test.

**Supplementary Table 1. Meteor localization sequences**

| Location | Terminus | Motif |
|---|---|---|
| cytoplasm | N | MLPPLERLTL |
| outer endoplasmic reticulum | N | MDPVVVLGLCLSCLLLLSLWKQSYGGGKL |
| mitochondrial matrix | N | MSVLTPLLLRGLTGSARRLPVPRAKIHSLGDP MSVLTPLLLRGLTGSARRLPVPRAKIHSLGDP KDPPVAT |
| nucleus | C | YEFPKKKRKVEDA |
